# A saturating mutagenesis CRISPR-Cas9 mediated functional genomic screen identifies *cis-* and *trans-* regulatory elements of *Oct4* in murine ESCs

**DOI:** 10.1101/851683

**Authors:** Matthew C. Canver, Pratibha Tripathi, Michael J. Bullen, Moshe Olshansky, Yogesh Kumar, Lee H. Wong, Stephen J. Turner, Samuel Lessard, Luca Pinello, Stuart H. Orkin, Partha Pratim Das

## Abstract

Regulatory elements (REs) consist of enhancers and promoters that occupy a significant portion of the non-coding genome and control gene expression programs either in *–cis* or in *– trans*. Putative REs have been identified largely based on their regulatory features (co-occupancy of ESC-specific transcription factors, enhancer histone marks and DNase hypersensitivity) in mouse embryonic stem cells (mESCs). However, less has been established regarding their regulatory functions in their native context. We deployed *cis-* and *trans-*regulatory elements scanning through saturating mutagenesis and sequencing (ctSCAN-SMS) to target elements within the ∼12kb *cis*-region (*Cis-*REs; CREs) of the *Oct4* gene locus, as well as genome-wide 2,613 high-confidence *trans-*REs (TREs), in mESCs. ctSCAN-SMS identified 10 CREs and 12 TREs, as novel candidate REs of the *Oct4* gene in mESCs. Furthermore, deletions of these candidate REs confirmed that the majority of the REs are functionally active, and CREs are more active than TREs in controlling *Oct4* gene expression. A subset of active CREs and TREs physically interact with the *Oct4* promoter to varying degrees; specifically, a greater number of active CREs compared to active TREs, physically interact with the *Oct4* promoter. Moreover, comparative genomics analysis reveals that more number of active CREs than active TREs are evolutionary conserved between mouse and primates, including human. Taken together, our study demonstrates the reliability and robustness of ctSCAN-SMS screening to identify critical REs, and investigate their roles in the regulation of transcriptional output of a target gene (in this case *Oct4)* in their native context.

## Introduction

Large-scale genomic studies reveal that ∼80% of the human genome may be involved in gene regulation, whereas only ∼2% of the genome encodes proteins (1). The functional non-coding genome can be broadly divided into regulatory elements (REs) and regions that encode non-coding RNAs (ncRNAs) (1, 2). Furthermore, REs can be sub-divided into *cis*-REs (CREs) and *trans*-REs (TREs), based on their position relative to their target gene(s). CREs are present proximally or distally relative to their target gene on the same chromosome, whereas TREs are located distally relative to their target gene on different chromosomes (3, 4). Putative REs have been identified using various methods, including transcription factor binding, enhancer histone marks, DNA accessibility (open chromatin regions), enhancer-promoter interactions, and gene expression (1, 5-10). REs usually enriched with sequence variants that are associated with diverse human traits and diseases (11-13). In addition, REs play crucial roles in evolutionary turnover and divergence (14-17).

Initial efforts systematically evaluated RE functions using reporter assays on a massive scale (18, 19); however, such approaches fail to interrogate REs functions within their native genomic contexts. Advances in CRISPR-Cas9 mediated genome editing technology (20, 21) have transformed analysis of protein-coding genes (22, 23), as well as REs *in situ* in chromatin. A few high-throughput CRISPR-Cas9 mediated functional genetic screens were performed to characterize CREs in mammalian cells (24-29). Prior screens to identify functional CREs were focused on – either targeting putative CREs of gene(s) of interest (gene-centric) or targeting putative CREs bound by selected TFs (TF-centric). Nevertheless, identification of functional TREs presents a challenge that has attracted less attention.

Here, we deployed genome-wide *cis-* and *trans-*regulatory elements scanning through saturating mutagenesis and sequencing (ctSCAN-SMS) in mouse embryonic stem cells (mESCs) to identify critical CREs and TREs of the *Pou5f1/Oct4* gene (a master pluripotency regulator of mESCs). We uncovered new functionally active CREs and TREs, and how they regulate *Pou5f1/Oct4* gene expression in mESCs.

## Results

### Design of a saturating CRISPR-Cas9 pooled library for ctSCAN-SMS

In mESCs, several putative REs, including 8,563 enhancers (ENs) and 231 super-enhancers (SEs) have been identified based on co-occupancy of ESC-specific TFs (OCT4, NANOG, SOX2, KLF4, ESRRB), mediators (MED1), enhancer histone marks (H3K4me1, H3K27ac) and DNase I hypersensitivity (30). SEs contain multiple ENs; SEs are also more densely co-occupied with TFs, enhancer histone marks, and chromatin regulators as compared to ENs, and associate with greater transcriptional output (30). We undertook a high-throughput CRISPR-Cas9 mediated genome editing approach to comprehensively target putative REs in mESCs. First, we generated a genome-wide map of open chromatin regions using ATAC-seq in mESCs, as ATAC-seq identifies most EN REs (10). ATAC-seq peaks were then overlayed within all putative ENs (8,563) and SEs (231) to designate “high-confidence” REs (2,613) (Fig. 1a; Fig. S1a; Table S1). As these REs are distributed genome-wide and on different chromosomes relative to *Oct4* gene locus (in *trans*-), we termed these REs as TREs. All possible single guide RNAs (sgRNAs) (20 nt) were designed (within the TREs for tilling) upstream of the *S. pyogenes* Cas9 NGG-protospacer adjacent motif (PAM) sequences to target the high-confidence 2,613 TREs (Fig.1a; Fig. S1a; Table S1 and S2). This analysis yielded 70,480 sgRNAs with a median gap of 5 bp between adjacent genomic cleavages (Fig. 1, c and d). Likewise, 1,827 sgRNAs were designed at the surrounding ∼12kb (−10kb to +2kb of TSS of the *Oct4*) region of the mouse *Oct4* gene locus to dissect the *cis-*REs (CREs) of *Oct4* (Fig. 1, b and c). In addition, the library included 2,000 non-targeting (NT) sgRNAs as negative controls, 119 sgRNAs targeting GFP (of the *Oct4*-GFP reporter that used for the screen) and 150 sgRNAs targeting coding sequence of mESC-TFs as positive controls (Fig. 1c). In total, the REs CRISPR-Cas9 pooled library contained 74,576 sgRNAs (Fig. 1c). These sgRNAs were synthesized, pooled, and cloned into a lentiviral vector for deep sequencing. The deep sequencing of the pooled library confirmed the presence of >95% sgRNAs that target TREs, >99% sgRNAs that target CREs and control sgRNAs (Fig. S1, b-f; Table S2).

**Figure. 1.**
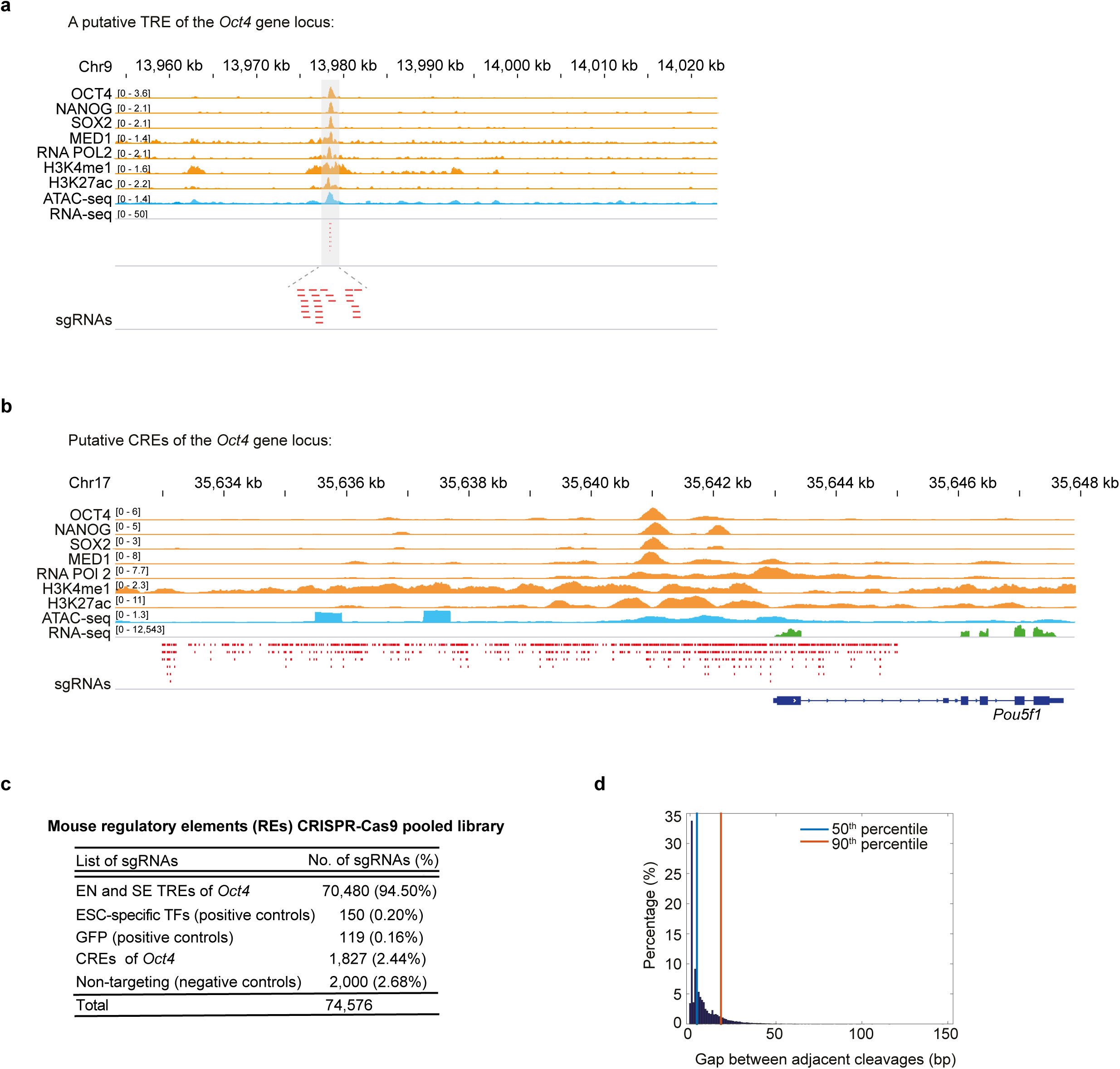
Design of a saturating CRISPR-Cas9 pooled library for ctSCAN-SMS. (a) Genomic tracks show co-occupancy of ESC-TFs (OCT4, NANOG, SOX2), mediator (MED1), enhancer histone marks (H3K27ac, H3K4me1), and RNA Pol2 at a putative TRE of the *Oct4* gene locus in mESCs. ATAC-seq track represents open chromatin regions and enhancers (ENs); RNA-seq track represents gene expression. Highlighted region displays as a putative TRE. All possible sgRNAs (shown with red dashed lines) were designed upstream of PAM sequences (NGG) within the putative TRE. (b) Genomic tracks illustrate co-occupancy of ESC-TFs (OCT4, NANOG, SOX2), mediator (MED1), enhancer histone marks (H3K27ac, H3K4me1) and RNA Pol2 at putative CREs of the *Oct4* gene locus in mESCs. ATAC-seq track characterises open chromatin regions and enhancers; RNA-seq track represents gene expression. sgRNAs (shown with red dashed lines) tilled upstream of PAM sequences (NGG) at the ∼12kb surrounding region (*-cis* region) of the *Oct4* locus. (c) Mouse REs CRISPR-Cas9 pooled library distribution. (D) Gaps between adjacent genomic cleavages of NGG PAM sgRNAs targeting CREs and TREs of the *Oct4*.

### Candidate CREs and TREs of the *Oct4* gene identified by ctSCAN-SMS

The pooled library was transduced into an *Oct4*-GFP reporter mESC line, which constitutively expresses Cas9 (31-33). The *Oct4*-GFP reporter was used as a “readout” for the screen to measure the reduction in GFP levels upon perturbation of any targeted RE regions by their corresponding sgRNAs. Lentiviral transduction of the pooled library was performed at low multiplicity (MOI) to ensure that each cell contained predominantly one sgRNA (Fig. S2a). After puromycin drug selection (since sgRNA constructs carrying puromycin drug resistance gene), “GFP-low” cells were sorted using fluorescence-activated cell sorting (FACS) (Fig. S2, a and b). As a control, cells were collected before FACS (the “pre-sort” sample). Genomic DNA was isolated from both “GFP-low” and “pre-sort” cell populations, and next-generation sequencing (NGS) was employed to enumerate the sgRNAs in each population (Fig. S2a). The screen was performed in triplicate.

We calculated an “enrichment score” of each sgRNA by comparing its frequency (presence) in GFP-low over pre-sort cells. Enrichment scores were determined based on the two best replicates (Table S2). Highest and lowest enrichment scores were obtained from GFP-targeting sgRNAs (mean log2FC 4.87, P-value <0.0001) and NT-sgRNAs (mean log2FC 0.44, P-value <0.0001), indicated that the screen was technically successful (Fig. 2a and 3a). We ranked all sgRNAs based on their enrichment scores (Table S2) and analyzed their off-target scores (ranged between 0-100) (34) (Fig. S3a and S4a; Table S2). A higher off-target score signifies fewer off-targets of a sgRNA. We found that the majority of the evaluated sgRNAs (87.6% sgRNAs for CREs, and 84.5% sgRNAs for TREs) had off-target scores >10 (Table S2); these sgRNAs are statistically significant based on the HMM (Hidden Markov Model) analysis (Fig. S3a).

**Figure. 2.**
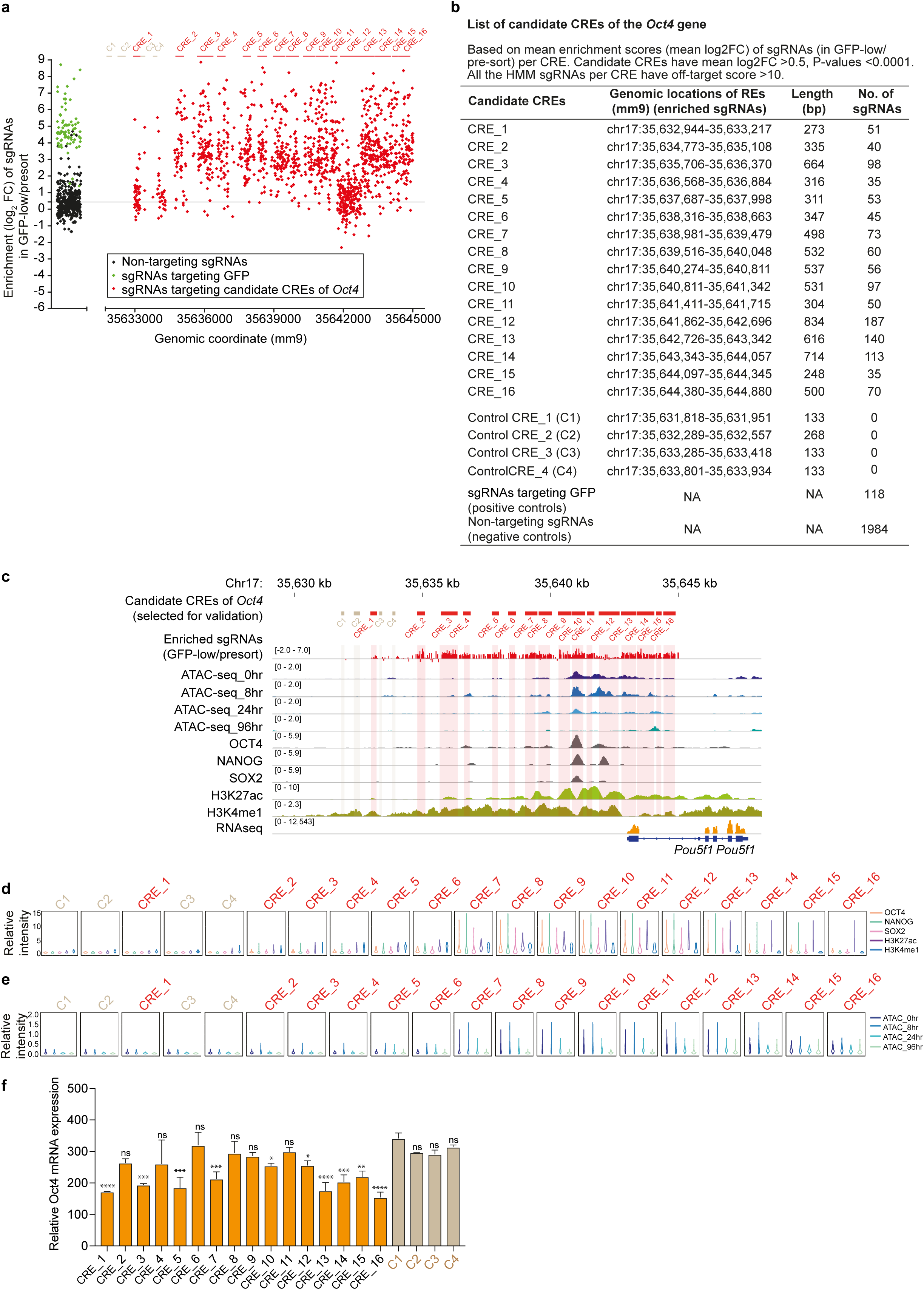
Identification and dissection of active CREs of the *Oct4* gene. (a) Dot plot analysis demonstrates the enrichment score of each sgRNAs by comparing their frequency in the GFP-low cells to the pre-sort cells. 16 candidate CREs were identified based on the mean enrichment score (mean log2FC) of sgRNAs per CRE. Four control CREs do not contain any sgRNAs. sgRNAs targeting GFP (green in color) and non-targeting sgRNAs (black in color) were used as positive and negative controls, respectively. (b) A list of identified candidate CREs of the *Oct4* gene. (c) Genomic tracks exhibit open chromatin regions/ENs by ATAC-seq at different time points (0, 8, 24, 96 hr) from undifferentiated to differentiated mESC state; co-occupancy of ESC-TFs (OCT4, NANOG, SOX2), enhancer histone marks (H3K27ac, H3K4me1) are also displayed at the mouse *Oct4* locus. RNA-seq shows the *Oct4* expression. (d) Violin plots outlining the binding dynamics of OCT4, NANOG, SOX2, H3K27ac, and H3K4me1 within the different CREs of *Oct4*. (e) Dynamic changes of open chromatin regions/ENs measured by ATAC-seq (using 0, 8, 24, 96 hr time points from undifferentiated to differentiated mESC state) within the CREs of *Oct4*. (f) Endogenous Oct4 mRNA expression levels quantified upon deletion of individual CREs of *Oct4*. Oct4 mRNA levels normalized to Gapdh. Data represented as mean ± SEM (n = 3); p-values calculated using ANOVA. *p < 0.05; **p < 0.01; ***p < 0.001; ****p < 0.0001; and ns (non-significant).

To identify candidate CREs of the *Oct4*, we considered all HMM sgRNAs with off-target scores >10 and mapped them within the ∼12kb surrounding region (−10kb to +2kb of TSS) of the *Oct4* gene locus. This yielded 16 candidate CREs (1-16), based on the mean enrichment score (mean log2FC) of sgRNAs per CRE. Each of the candidate CREs had a mean log2FC>0.5 (range of mean log2FC 0.93-3.95), with P-values <0.0001, which was higher than the mean enrichment score of non-targeting (NT)-sgRNAs (mean log2FC 0.44, with P-value <0.0001) (Fig. 2, a and b). Among 16 CREs, CREs-10 and 12 have been recognized previously as distal and proximal enhancers, respectively (31); CREs-13 to 16 were present within the promoter region of *Oct4* (+/-2kb of TSS) (31). The remaining 10 CREs were newly identified candidate CREs of the *Oct4* gene (Fig. 2, a and b). Of note, CRE-12 displayed a relatively lower mean enrichment score (log2FC 0.77, P-values <0.0001) compared to others candidate CREs, because the original *Oct4*-GFP reporter lacks a portion of CRE-12 (33). Nonetheless, we included CRE-12 for further validation.

To classify the candidate TREs, we applied a statistically relevant Hidden Markov Model (HMM) to the sgRNAs enrichment scores (35), which initially identified 263 candidate TREs. Furthermore, we applied stringent criteria to select candidate TREs for validation, as follows: i) TREs must have HMM sgRNAs with off-target scores>10 (Fig. 3a and Fig. S4a); ii) TREs must possess at least 4 sgRNAs with mean log2FC >0.5 (range of mean log2FC 0.86-2.52), with P-values <0.0001 (Fig. 3b); iii) TREs must co-occupy with ESC-TFs (OCT4, NANOG, SOX2), enhancer histone marks (H3K27ac, H3K4me1) (Fig. 3c and Fig. S4b); and iv) they must contain “dynamic” open chromatin regions; i.e. “open” chromatin regions present in the undifferentiated state (0 hr) that gradually become “closed” with differentiation (24, 96hr) of mESCs (Fig. 3d and Fig. S4c). Based on these criteria, we selected 12 candidate TREs of the *Oct4* gene (Fig. 3, a and b).

**Figure. 3.**
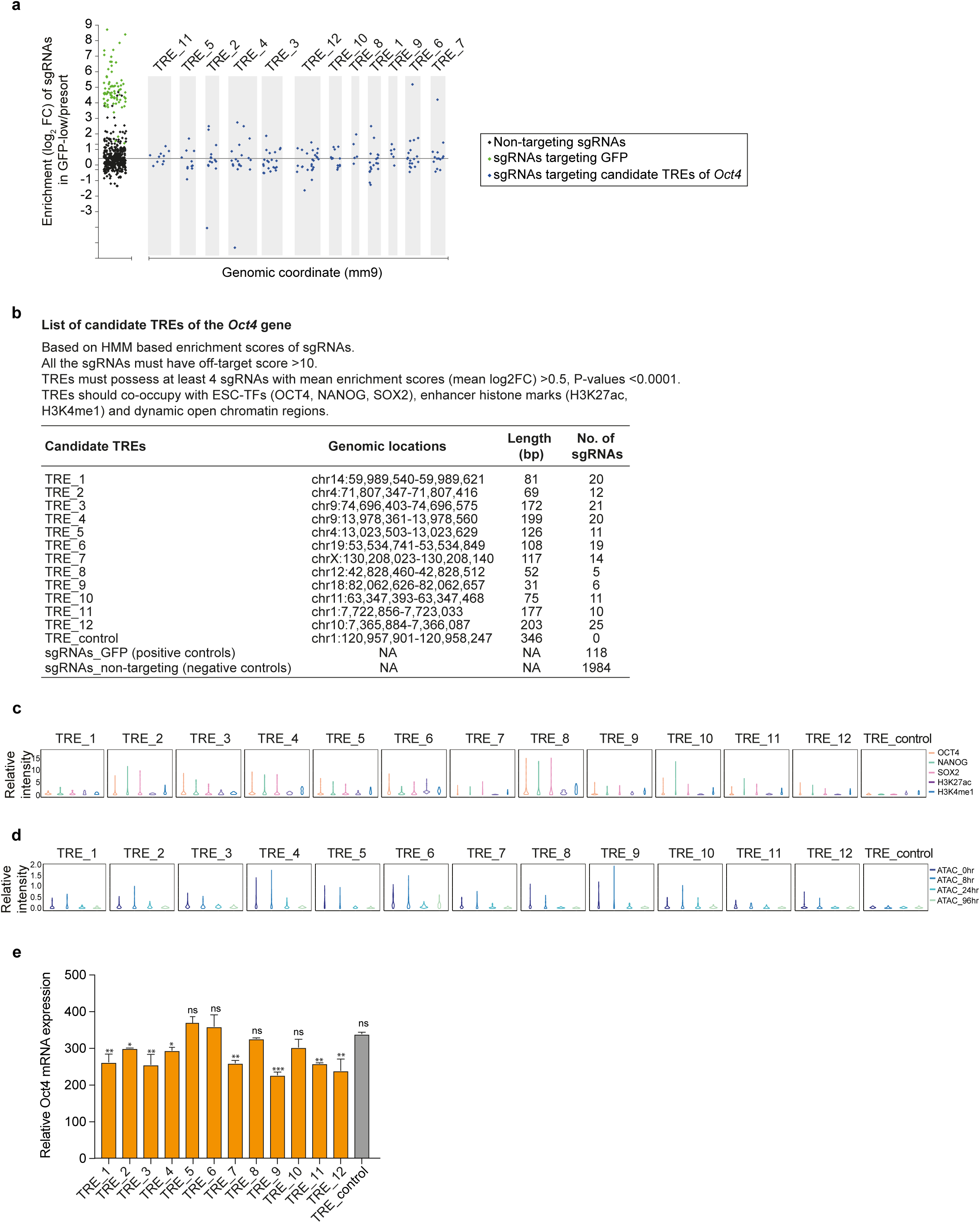
Identification and validation of active TREs of the *Oct4* gene. (a) Dot plot analysis displays the enrichment score of each sgRNAs by comparing their frequency in the GFP-low cells to the pre-sort cells at the selected candidate 12 TREs. (b) A list of identified candidate TREs of the *Oct4* gene. (c) Binding dynamics of OCT4, NANOG, SOX2, H3K27ac, and H3K4me1 represented within the TREs of *Oct4*. (d) ATAC-seq demonstrates the changes in open chromatin regions/ENs from undifferentiated to differentiated mESC state (0, 8, 24 and 96 hr) within the TREs. (e) Quantitative RT-PCR data illustrates the mRNA expression changes of endogenous Oct4 upon deletion of individual TREs of *Oct4*. Oct4 mRNA levels normalized to Gapdh. Data represented as mean ± SEM (n = 3); p-values calculated using ANOVA. *p < 0.05; **p < 0.01; ***p < 0.001; and ns (non-significant).

### Dissection of functionally active CREs and TREs of the *Oct4* gene

We selected a total of 33 REs, including 20 CREs – 16 candidate CREs and 4 control CREs (CREs with no significant enrichment scores of sgRNAs), and 13 TREs – 12 candidate TREs and one control TRE (with no significant enrichment scores of sgRNAs) of the *Oct4* gene locus for validation using wild-type mESCs. “Paired” sgRNAs (5’ and 3’ sgRNAs) tagged with mCherry were used to target both the flanking ends of each selected candidate RE to create a deletion. Following transfection of paired sgRNAs-mCherry into the wild-type mESCs, mCherry-positive cells were sorted to measure the endogenous Oct4 mRNA expression and to confirm the genomic deletions of CREs/TREs (Fig. 2f and 3e; Fig. S3d; Fig. S4e; Table S3). We observed a significant reduction in Oct4 expression to different extents upon deletions of CREs-1, 3, 5, 7, 10, 12, and 13-16 (out of 16 candidate CREs) (Fig. 2f). Deletions of newly identified CREs-1, 3, 5 and 7 showed a greater reduction in Oct4 expression, compared to deletions of known distal and proximal enhancers (CREs-10 and 12) of *Oct4* (Fig. 2f). However, “regulatory features”, such as co-occupancy of ESC-TFs (OCT4, NANOG, SOX2– ONS), enhancer histone marks (H3K27ac, H3K4me1) and dynamic open chromatin regions (ATAC-seq peaks at 0hr compared to 24, 96hr), were more prominent at CREs-7, 10, 12 compared to CREs-1, 3, 5 (Fig. 2, c-e; Fig. S3, b and c). Moreover, deletions of CREs-13 to 16 (present at the promoter region of *Oct4*) showed a significant reduction in Oct4 expression, as expected (Fig. 2f). Nonetheless, only CREs-13, 14 showed substantial co-occupancy of ONS, H3K27ac and dynamic open chromatin regions, as compared to other CREs present at the *Oct4* promoter (Fig. 2, c-e; Fig. S3, b and c). In contrast, deletions of control CREs (C1-C4) led to no significant changes in Oct4 expression (Fig. 2f). Additionally, control CREs exhibited low-level co-occupancy of ONS, H3K27ac, H3K4me1, and dynamic open chromatin regions (Fig. 2, d and e; Fig. S3, b and c). These data confirm the existence of multiple “active” CREs (“active” REs depict – REs that reduce the Oct4 expression, upon their deletions), including newly identified active CREs of the *Oct4*. Yet some “active” CREs fail to display recognized regulatory features (i.e. without of any co-occupancy of TFs, enhancer histone marks, and open chromatin regions) as described recently (26, 28), suggesting that a subset of functionally “active” REs may lack typical regulatory features.

Deletions of 8 TREs (TREs-1, 2, 3, 4, 7, 9, 11, 12) of 12 candidates showed a significant reduction in Oct4 expression to various extents (Fig. 3e). Nonetheless, all 12 TREs were co-occupied with ONS, H3K27ac and H3K4me1 enhancer marks, and exhibited dynamic open chromatin regions (Fig. 3, c and d; Fig. S4, b and c); as all the candidate TREs (TREs 1-12) were short-listed for validation based on their regulatory features. Conversely, control TRE lack typical regulatory features (Fig. 3, c and d); and deletion did not affect Oct4 expression (Fig. 3e). Moreover, neighbouring genes of most of the validated active TREs were lowly expressed in mESCs (Fig. S4d), indicating that these genes may not be critical for the mESC state maintenance. Importantly, we observed reduction of Oct4 expression was greater for the majority of active CREs than active TREs, upon their deletions (Fig. 2f and 3e). This suggests that CREs generally contribute more than TREs to control overall *Oct4* gene expression. Taken together, these data demonstrate that reliability and robustness of the ctSCAN-SMS screen to identify new candidate REs of the *Oct4* in a high-throughput manner. Moreover, we confirm that a subset of these newly identified candidate CREs and TREs are functionally “active” REs of the *Oct4* gene, and CREs are more active compared to TREs.

### *Cis-* and *trans-* regulation of the *Oct4* gene expression

REs (particularly enhancers) physically interact with gene promoters and control transcription (36). Chromosome conformation capture (3C) based methods – 4C, Hi-C, capture Hi-C, ChiA-PET and HiChIP--have been utilized to identify physical contacts between promoters and REs (enhancers) (4). To interrogate the potential mechanisms by which candidate CREs and TREs regulate *Oct4* gene expression, we examined interactions between REs (CREs and TREs) and the *Oct4* promoter using published 4C-seq data. These data were generated to study intra-chromosomal and inter-chromosomal interactions between REs and the *Oct4* promoter at a genome-wide scale (37). We used Oct4-234 (a region at ∼1.5kb upstream of TSS of *Oct4*) as a “viewpoint”, as previously (Fig. S5a) (37). Next, “contact frequencies” were calculated between the viewpoint and CREs (using 1kb resolution window, surrounding 30kb region of the *Oct4* gene locus) (Fig. 4, a and c), as well as between the viewpoint and TREs (using 50kb resolution window, surrounding each of the TREs) (Fig. 4b). This analysis revealed ranges of contact frequencies between functionally validated “active” CREs/TREs and the *Oct4* promoter (Fig. 2f and 3e; Fig. 4, a and b). For example, functionally validated “active” CREs, including CREs-3, 5, 7 (newly identified CREs), CREs-10 and 12 (previously known as distal and proximal enhancers of *Oct4*), and CREs-13 to16 (residing at the promoter region of *Oct4*) demonstrated intra-chromosomal interactions with the *Oct4* promoter (Fig. 4, a and c). Established enhancers, such as CREs-10 and 12, showed higher contact frequencies compared to newly identified CREs-3, 5, 7 (Fig. 4, a and c). However, active CRE-1 and control CREs (C1-C4) did not show significant interactions with the *Oct4* promoter (Fig. 4, a and c). In contrast, among functionally validated “active” TREs-1, 2, 3, 4, 7, 9, 11, 12; only a minority (TREs-2, 3, 4, 11) displayed higher inter-chromosomal interactions (compared to control TRE) with the *Oct4* promoter (Fig. 4b).

**Figure. 4.**
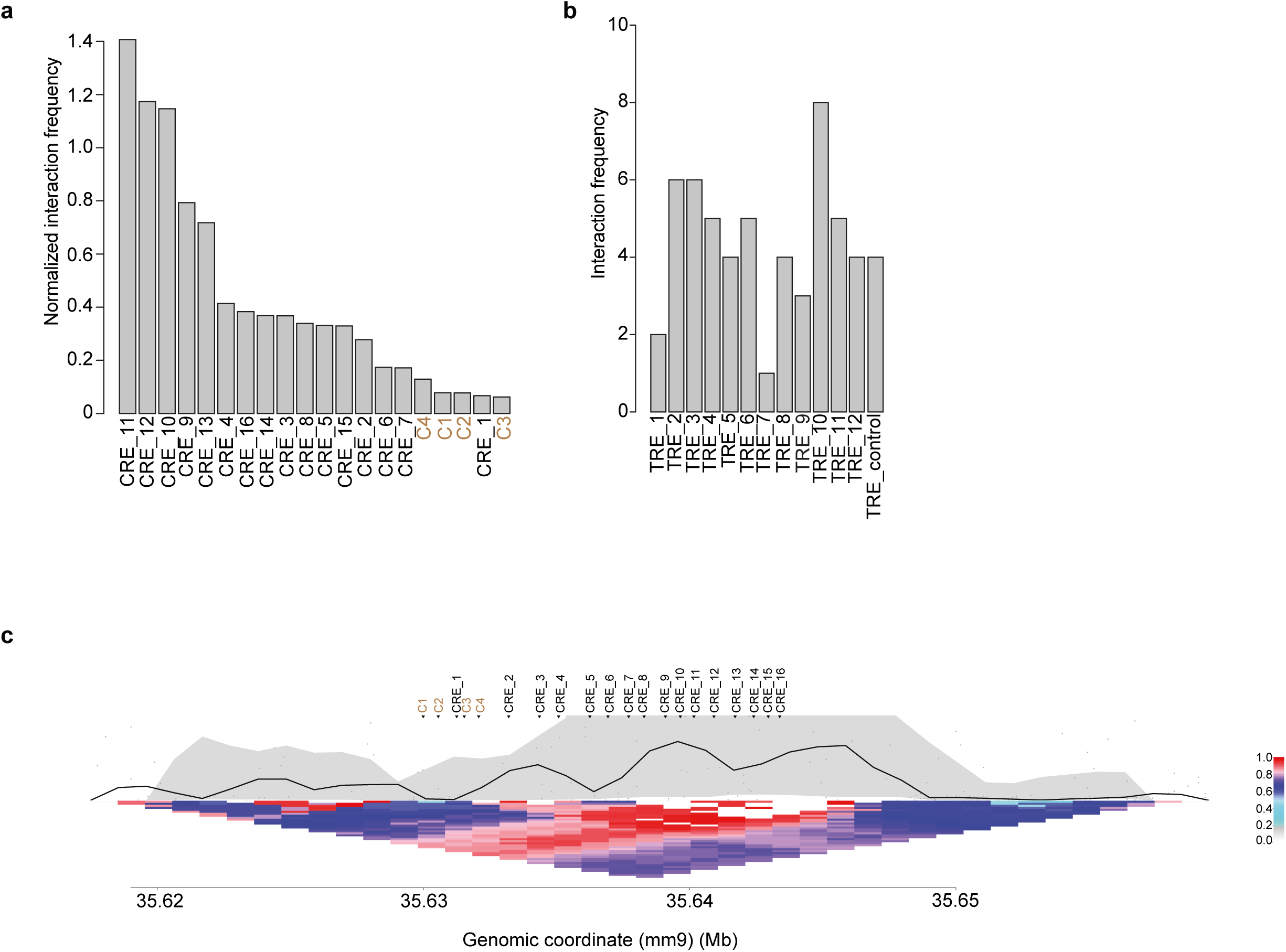
Physical interactions between CREs, TREs, and the *Oct4* promoter in *Oct4* gene regulation. (a) 4C-seq data represent normalized interaction frequencies between CREs and the *Oct4* promoter. The interaction/contact frequencies between CREs and the *Oct4* promoter measured at a 1kb resolution window. (b) The interaction/contact frequencies between TREs and the *Oct4* promoter quantified at a 50 kb resolution window. (c) The contact profile of CREs and the *Oct4* promoter at a 1kb resolution window. Bottom triangle is a heat map of normalized contact frequencies between 1kb bins represented with the color codes (the highest is 1 with red; the lowest is 0 with white in color). At the upper part, the black line (within the grey region) represents the normalized median contact frequencies between a locus and the viewpoint. Grey region exhibits 20^th^-80^th^ percentile of the normalized contact frequencies.

Likewise, a high-resolution Micro-C (Micrococcal nuclease (MNase) based Hi-C assay that captures genome-wide 3D chromatin organization/contact frequencies at single nucleosome resolution (∼100-200bp)) from mESCs (38), displayed ranges of intra-chromosomal interactions between active CREs and the *Oct4* promoter (Fig. S5, b and c). Further, these data evaluated all other intra-chromosomal interactions between any two genomic loci around ∼20kb region of the *Oct4* gene locus at 200bp resolution (Fig. S5b). In addition, inter-chromosomal interactions were also observed between a few active TREs (TREs-4, 11) and the *Oct4* promoter (Fig. S5d). Both 4C-seq and Micro-C data showed that greater contact frequencies of active CREs compared to active TREs with the *Oct4* promoter, and a greater number of active CREs compared to active TREs physically interacted with the *Oct4* promoter (Fig. 4, a-c; Fig. S5, b-d).

Taken together, our data suggest that a subset of active CREs and TREs physically interact with the *Oct4* promoter at different frequencies as they influence the *Oct4* transcriptional output.

### Conserved functionally active CREs and TREs of the *Oct4* gene

Recent studies demonstrate that the majority of species-specific REs/ ENs evolved *de novo* from ancestral DNA regulatory sequences (16, 39). Also, evidence implies that loss or gain of REs (called RE turnover) takes place during evolution (17). To understand the importance of validated active CREs and TREs of the mouse *Oct4* gene in evolutionary turnover, we analyzed their regulatory sequence conservation. Active CREs-3, 5; CREs-10 and 12 (known distal and proximal ENs); CREs-13 to 16 (present within the promoter) demonstrated significant conservation between mouse and primates, including human (Fig. 5a); whereas active CREs-1, 7 did not show appreciable sequence conservation between mouse and primates (Fig. 5a). In comparison to active CREs, only a small subset of active TREs (TREs-3 and 4) showed significant sequence conservation between mouse and primates (Fig. 5, b and c). These observations suggest that active CREs are more critical than active TREs for *Oct4* expression in mouse and primates (including human) during evolution.

**Figure. 5.**
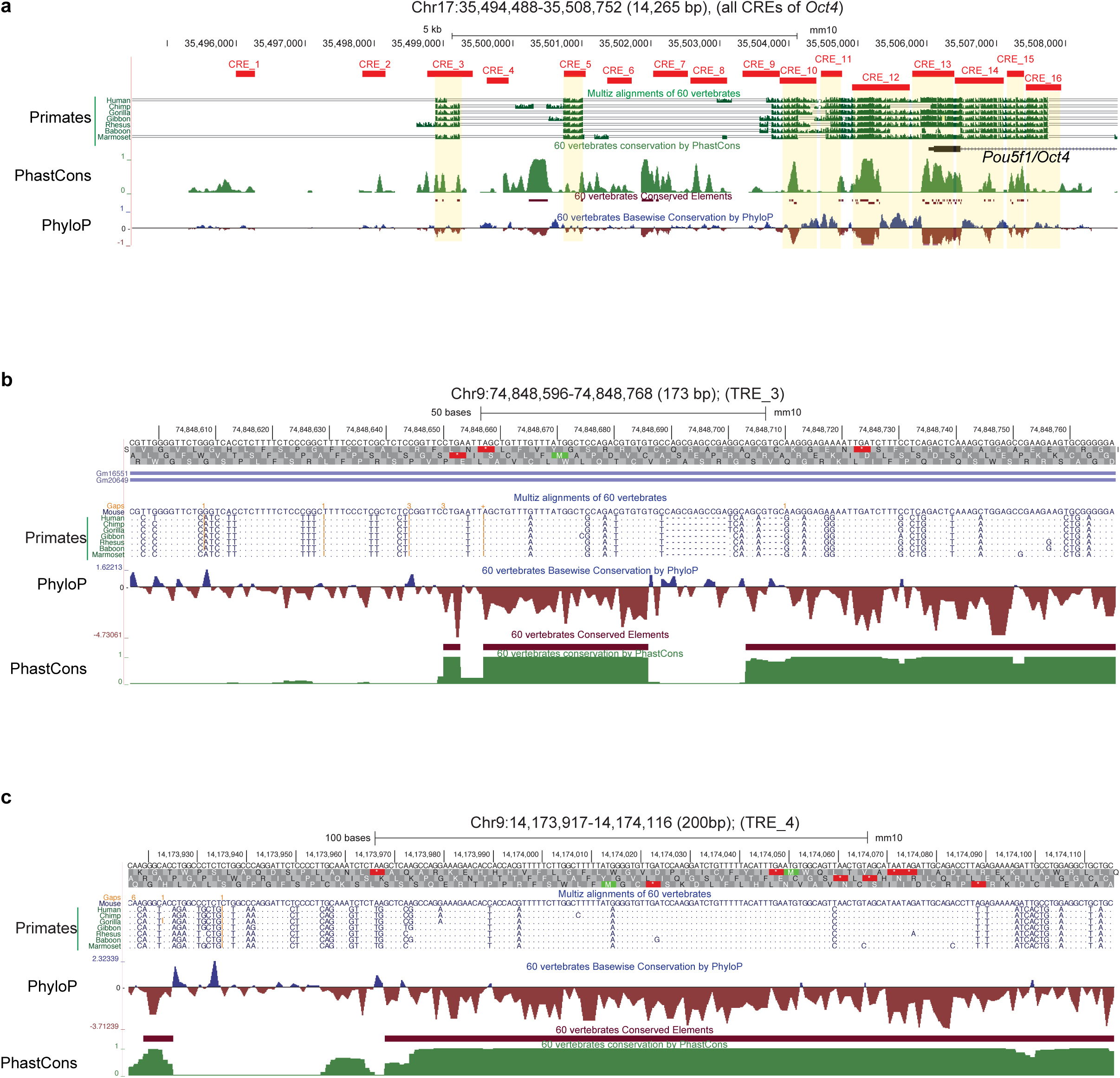
Conserved active CREs and TREs of the *Oct4* gene. (a) Orthologous sequences from the representative primates (including human) are listed around the ∼12kb region of the mouse *Oct4* locus. CREs of the mouse *Oct4* are labelled with solid red bars. PhyloP and PhastCons estimate evolutionary conservation among 60 vertebrates. (b-c) Orthologous sequences from the representative primates (including human) are listed at the active TRE-3 (b) and TRE-4 (c) regions of the mouse *Oct4*. PhyloP and PhastCons estimate evolutionary conservation among 60 vertebrates.

Next, we analyzed regulatory sequence conservation of previously identified high-confidence CREs (−449, -571, -694) of human *OCT4*. These CREs are located distally between ∼450 to 700kb upstream of the human *OCT4* TSS, and physically interact with the *OCT4* gene (28). We found that these human CREs are also evolutionary conserved at the upstream regions of the mouse *Oct4* locus (Fig. S6).

Taken together, our comprehensive comparative genomic analysis shows that a larger subset of active CREs compared to active TREs of *Oct4* are well-conserved between mouse and primates including human. These observations support the existence of both conserved, as well as non-conserved REs of *Oct4* among mouse and primates (including human), which is consistent with “RE turnover/ divergence” (gain and loss of REs) at the *Oct4* locus during evolution that may be critical for positive selection, as proposed earlier (17, 39).

## Discussion

A handful of modest scale CRISPR-Cas9 mediated functional screens (using hundreds to thousands of sgRNAs) have been performed to target specific non-coding CREs of gene(s) of interest (24-26, 29, 40). These screens successfully identified functional CREs of the target gene(s). In the context of identification of CREs of the *OCT4* gene, a previous CRISPR-Cas9 mediated screen was performed to target 174 candidate CREs of *OCT4* within its 1MB topological associated domain (TAD) in human ESCs; it revealed 4 temporary CREs and 2 known proximal CREs. The temporary CREs showed “transient” enhancer regulatory activity in *OCT4* gene expression (27). However, the functional relevance of these temporary CREs is uncertain in human *OCT4* gene regulation. Furthermore, another CRISPR-Cas9 mediated screen was performed by the same group using a different strategy, called CREST-seq. This method employed 11,570 paired sgRNAs to introduce deletions to target 2Mb surrounding the *OCT4* locus in human ESCs (hESCs), which created 2kb deletions on average with an overlap of 1.9kb between two adjacent deletions. This screen identified total 45 CREs, of which 17 CREs (with regulatory features) reside at the promoters of “unrelated” genes (intra-chromosomally) that act as typical enhancers of the *OCT4* gene (28). Our study employed ctSCAN-SMS – an unbiased, high-resolution, high-throughput screening approach using 1,827 sgRNAs to target CREs and 70,480 sgRNAs to target TREs of the mouse *Oct4* gene (Fig. 1).

Previous CRISPR-Cas9 screens were focused on CREs of the target gene(s). In contrast, our screen was designed to identify both CREs and TREs. Indeed, we discovered 16 CREs (including 10 novel CREs) and 12 novel TREs of the murine *Oct4* gene, as potential candidate REs (Fig. 2a and 3a). Furthermore, deletion studies confirmed that the majority of these CREs (10 CREs out of 16 CREs) and TREs (8 TREs out of 12 TREs) are functionally “active” in controlling the *Oct4* expression. Overall, CREs are more functionally active than TREs (Fig. 2f and 3e). In addition, we showed that a subset of these active CREs and TREs physically interacts with the *Oct4* promoter to different extents through intra- and inter-chromosomal interactions, respectively (Fig. 4; Fig. S5). Notably, a greater number of active CREs, compared with active TREs, physically interact with the *Oct4* promoter. Moreover, the interactions between active CREs (compared to active TREs) and the *Oct4* promoter are more prominent (Fig. 4, Fig. S5). Nonetheless, “enhancer activities” of several active CREs/TREs are not directly correlated to their physical contact frequencies with the *Oct4* promoter (Fig. 2f, 3e; Fig. 4; Fig. S5). Interestingly, we found several active CREs (CREs-1, 3, 5) lack typical regulatory features (Fig. 2, d-f; Fig. 6); this supports earlier studies that identified unmarked REs (UREs) without typical regulatory features may play critical roles in transcriptional output (26, 28). Moreover, comparative genomics analysis revealed that numerous active CREs and active TREs (more number of active CREs than active TREs) of *Oct4* are evolutionarily conserved between mouse and primates (including human) (Fig. 5). However, we also observed divergence of *Oct4* REs among mouse and primates (including human), which may account for the vital roles of “RE turnover” during evolution (3, 39).

**Figure. 6.**
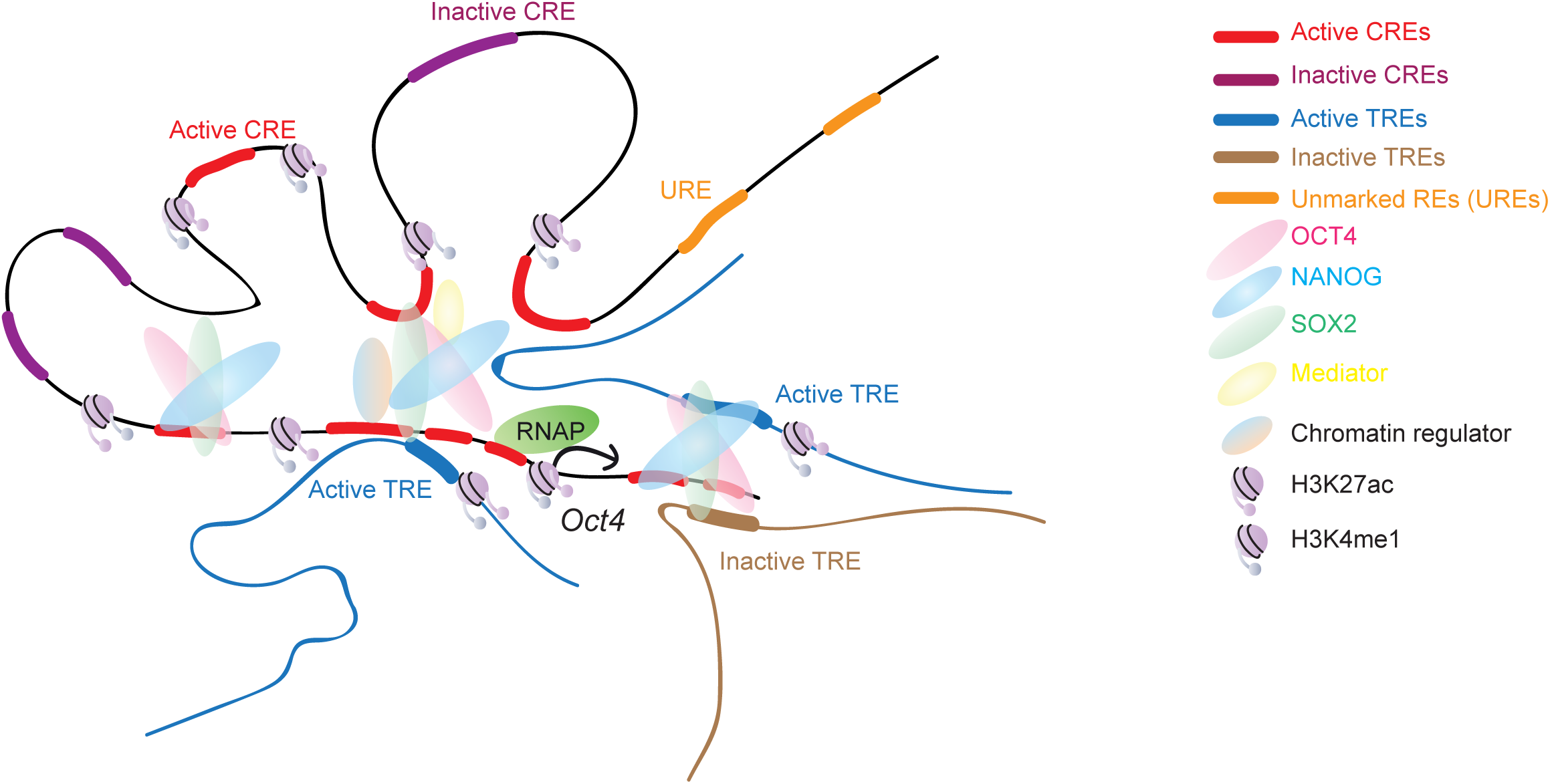
Model representing the detailed functions of CREs and TREs in *Oct4* gene regulation. The proposed model describes the existence of multiple “active” CREs (red in color) and TREs (blue in color) of the *Oct4* locus in mESCs. However, not all of the active REs have regulatory features (i.e. co-occupancy of ESC-TFs (OCT4, NANOG, SOX2), active enhancer histone marks (H3K27ac, H3K4me1), and open chromatin regions), which are termed as unmarked REs (UREs). Also, a subset of active REs physically interacts with the *Oct4* promoter through intra-chromosomal (for CREs) and inter-chromosomal (for TREs) interactions. Taken together, it suggests that active REs act beyond their regulatory features and physically contact with the *Oct4* promoter to control the transcriptional output of *Oct4* gene.

Several studies demonstrate that “multiple REs” act either in a co-operative or competitive fashion to control transcriptional output (39). Our study identified multiple active REs of *Oct4* and revealed a spectrum of regulatory activities of “individual” CREs and TREs in *Oct4* gene expression (Fig. 2f and 3e). Further systematic studies will be required to elucidate how multiple REs function combinatorially to control *Oct4* transcriptional output. In conclusion, we have demonstrated the utility of ctSCAN-SMS as an approach to identify functional REs of a gene locus, and dissect their regulatory contributions to the transcriptional output of a target gene within its normal chromosomal context.

## Experimental procedures

### Mouse embryonic stem cells (mESCs)

Mouse ESCs (mESCs) were cultured in mouse ESC media that contains DMEM (Dulbecco’s modified Eagle’s medium) (Thermo Fisher Scientific) supplemented with 15% fetal calf serum (FCS) (Merck Millipore), 0.1mM b-mercaptoethanol (Sigma-Aldrich), 2mM L-glutamine (Thermo Fisher Scientific), 0.1mM nonessential amino acid (Thermo Fisher Scientific), 1% of nucleoside mix (Merck Millipore), 1000 U/ml recombinant leukemia inhibitory factor (LIF/ESGRO) (Merck Millipore), and 50U/ml Penicillin/Streptomycin (Thermo Fisher Scientific), as described previously (32, 41). mESCs were cultured at 37°C, 5% CO_2_.

### Mouse REs CRISPR-Cas9 pooled library design for the ctSCAN-SMS

For this study, we selected a list of putative 8,563 enhancer (EN) and 231 super-enhancer (SE) REs from the mESCs, as described previously (30). First, we generated ATAC-seq (10) data from wild-type mESCs, and mapped within all the putative EN and SE REs to identify open chromatin regions, as well as high-confidence REs. Next, ±100 bp (200 bp) around the centre of the ATAC-seq peaks were obtained from the high-confidence REs. In total, we identified 2,613 REs for targeting. All possible single guide RNAs (sgRNAs) (20nt) were designed upstream of the *S. pyogenes* Cas9 NGG-protospacer adjacent motif (PAM) sequences at these defined REs (2,613), which created 70,480 sgRNAs with a median gap 5 bp between adjacent genomic cleavages. Since these EN and SE REs distributed in *trans-* of the *Oct4* gene locus, we called these REs as *trans-*REs (TREs). Likewise, 1,827 sgRNAs were designed prior to all possible NGG-PAM sequences at the adjacent ∼12kb (−10kb to +2kb of TSS of *Oct4*) region of the mouse *Oct4* gene locus to systematically dissect the REs of *Oct4*. As these REs reside adjacent to the *Oct4* gene locus, they are called *cis-*REs (CREs). We also included 2,000 non-targeting (NT) sgRNAs as negative controls; 119 sgRNAs targeting GFP of the *Oct4*-GFP reporter and 150 sgRNAs targeting coding sequence of mESC-TFs as positive controls. Altogether, the REs CRISPR-Cas9 pooled library contained total 74,576 sgRNAs.

### REs CRISPR-Cas9 pooled library construction for the ctSCAN-SMS

All the sgRNA oligonucleotides of the library were synthesized as previously described (32) using a B3 synthesizer (CustomArray, Inc.), pooled together, PCR amplified and cloned into Esp3I-digested plentiGuide-Puro (Addgene plasmid ID: 52963) lentiviral vector, using a Gibson assembly master mix (New England Biolabs).Gibson assembly products were transformed into electrocompetent cells (*E. cloni*, Lucigen) and plated on 245mm x 245mm square LB-agar plates to obtain the sufficient number of bacterial colonies at a ∼50× library coverage. Bacterial colonies were collected from the plates, genomic DNA was isolated and plasmid libraries were prepared for high-throughput sequencing to confirm the representation of individual sgRNA in the REs CRISPR-Cas9 pooled library.

### Lentiviral library production

HEK293T cells were seeded onto 15cm dishes ∼24hrs prior to transfection. Cells were transfected at 80% confluence in 16ml of media with 8.75μg of VSVG, 16.25μg of psPAX2, and 25μg of the REs CRISPR-Cas9 pooled lentiviral plasmids, using 150μg of linear polyethylenimine (PEI) (Sigma-Aldrich). Media was changed with fresh media, 16–24hrs after transfection. Lentiviral supernatant was collected at 48 and 72hrs post-transfection and subsequently concentrated by ultracentrifugation (24,000 rpm, 4°C, 2hrs) (Beckman Coulter SW32).

### CRISPR-Cas9 mediated ctSCAN-SMS in mESCs

*Oct4*-GFP reporter mESCs with stably expressed Cas9 were transduced with REs CRISPR-Cas9 pooled lentiviral library at low multiplicity of infection (MOI) to avoid more than one lentiviral integration per cell. Test transductions were performed to estimate the viral titration and transduction rate. Briefly, 300,000 *Oct4*-GFP+Cas9 mESCs were plated per well of a 12-well plate. After 24hrs, different amounts of (1, 2, 4, 6, 8µl) of the lentiviral library was added to the cells. 10μg/ml blasticidin (InvivoGen) and 1μg/ml puromycin (Sigma-Aldrich) were added 24hrs after the transduction to select for lentiviral library integrants (puromycin resistant) in cells with Cas9 (blasticidin resistant). Cells were selected for the next 3-4 days. The same number of cells were seeded as a control; but not infected with lentiviral library and not treated with blasticidin and puromycin. The number of blasticidin and puromycin resistant cells, and control cells were counted to calculate the viral titre and transduction rate (to achieve 30%).

For the actual screen, we seeded ∼112 million (∼75K sgRNAs in the pooled library, with 500X coverage, for 30% transduction rate) *Oct4*-GFP+Cas9 mESCs in the same format (i.e. 300K cells/ well of the 12-well plate) for each independent screening replicate. Lentiviral library was added to each well of 12-well plate to achieve 30% transduction rate with low MOI (MOI 0.1) to make sure each infected cell obtained one viral particle. 24hrs post-transduction, fresh mESC media was added to the cells with 10μg/ml blasticidin (InvivoGen) and 1μg/ml puromycin (Sigma-Aldrich) and selected for 4 days. These selected cells were used to sort the “GFP-low” cells. The “pre-sort” cells were collected before sorting and used as a control. Genomic DNA was isolated from both the “GFP-low” and “pre-sort” cell populations, libraries were prepared for deep sequencing to enumerate the presence of sgRNAs in these cell populations as previously described (32). The screening was performed in biological triplicates.

### Deletion of CREs and TREs of *Oct4* in mESCs

Paired sgRNAs (5’ and 3’ sgRNAs) were designed to target both the ends of each selected candidate RE to create a deletion. Both the sgRNAs were cloned into lentiguide-plasmid that carries Cas9 and mCherry, using Golden Gate Cloning approach as previously described (32). 1,000,000 wild-type (J1) mESCs were transfected with 1µg of each 5’ and 3’ sgRNA-Cas9-mCherry plasmids using Lipofectamine 2000 (Thermo Fisher Scientific). After 24hrs of transfection, mCherry-positive cells were sorted; at least 50,000-100,000 mCherry-positive cells were collected either to isolate the total RNA for measuring the endogenous Oct4 mRNA expression levels by quantitative RT-PCR or to perform the genotyping PCR for checking deletions of targeted REs.

### Flow cytometry

Cells were dissociated using trypsin, washed with 1XPBS, followed by sorting. i) During the CRISPR-Cas9 screening, *Oct4*-GFP reporter mESCs were sorted based on GFP-low intensity after the transduction with REs CRISPR-Cas9 pooled library; ii) to quantify the Oct4 mRNA expressions upon deletions of CREs and TREs, their targeting sgRNAs-mCherry-positive cells were sorted.

### RNA isolation and RT-qPCR

DNA-free total RNA was isolated from mESCs using RNeasy Mini Kit (Qiagen), and cDNA was prepared using iScript cDNA Synthesis Kit (Bio-Rad). RT-qPCR was performed using iQ SYBR Green Supermix (Bio-Rad) on Bio-Rad iCycler RT-PCR detection system.

### ctSCAN-SMS screen data analysis

sgRNA sequences present in the GFP-low and pre-sort pools were enumerated. Enrichment was determined by the log2 transformation of the median number of occurrences of a particular sgRNA in the GFP-low pool divided by the median number of occurrences of the same sgRNA in the pre-sort pool across the best two biological screen replicates.

### ATAC-seq experiment and data analysis

ATAC-seq was performed according to the previously described protocol (10), with some modifications. Briefly, each library was started with 50,000 cells, which were washed with 1X PBS and permeabilized with 50µl of lysis buffer (10mM Tris, pH 7.4; 10mM NaCl; 3mM MgCl2; 0.1% IGEPAL) at 4°C by resuspension. Cells were centrifuged at 500g for 10 min at 4°C to pellet the nuclei. The resulting nuclei were resuspended in 50µl of transposition reaction buffer (25µl of 2x TD buffer from the Nextera kit, Illumina; 2.5µl of Tn5 transposase enzyme from the Nextera kit, Illumina; 22.5µl of nuclease free water), and incubated at 37°C for 90 min for chromatin tagmentation. Next, DNA was purified using Qiagen MinElute PCR purification kit, and eluted in 10µl of nuclease-free water. PCR amplification was performed using Nextera primers (Illumina) to make the libraries for deep sequencing.

The obtained deep sequencing data in .FASTQ format was inspected first by FASTQC. Next, reads were trimmed for adapters using Trimmomatic (42). The resulting fastq files were aligned with Bowtie2 (43) with the following options --local -X 2000. Peaks were called with MACS2 (44) with the following options callpeak --gsize mm --nomodel --shift -100 --extsize 200 --call-summit.

### 4C-seq data analysis

The normalised interaction frequencies between CREs and *Oct4* promoter measured based on 4C-seq pipeline (37), with some modification. Normalised interaction frequencies between CREs and *Oct4* promoter (Oct-234 used as a view point) was quantified at a higher resolution (1kb resolution window compared to previous analysis used 7kb resolution window). We used the following command:

perl <u>4cseqpipe.pl</u> -dopipe -ids 1 -fastq_fn Oct4/fastq/Oct4_234.fastq -convert_qual 1 - calc_from 35620000 -calc_to 35660000 -stat_type median -trend_resolution 1000 -figure_fn Oct4_234_1K.pdf -feat_tab rawdata/Oct4_234_features.txt

The contact frequencies between TREs and *Oct4* promoter was calculated based on the number of contacts between the *Oct4* promoter (Oct4-234 used as a view point) and a 50kb region centred at each TRE, using bedtools intersect command.

### Micro-C data analysis

Raw Micro-C data from mouse ESC cells (mESCs) was downloaded from GEO: GSE130275 (38). Samples were processed with a standard juicer pipeline (45), using mm9 mouse genome. Reads from all 37 samples were merged together (after duplicates removal). This resulted in more than 4 billion contacts, allowing the creation of Hi-C file at a very high (200 bp) resolution. For CRE contacts, the chr17 intra-chromosomal contacts map was processed using in house tool to produce normalized observed/expected counts. These counts were used for the creation of heatmap/matrix and barplot. For each CRE (including *Oct4* TSS), all 200bp bins overlapping the loci were used. For TRE contacts, whole-genome 1kb resolution contacts map was created and balanced using our in house C++ code (this is the GW_SCALE normalization available in juicer_tools_1.22.01.jar – *manuscript in preparation*). These counts were used for the creation of the barplot. In this case, *Oct4* promotor was defined as -/+ 3kb of TSS, and TREs were used 200kb bin centered at the particular TRE.

The heatmap was generated using a heatmap.2 R function from gplots package and barplots were generated using a barplot R function.

## Data availability

All the high-throughput sequencing data from this study have been submitted to the NCBI Gene Expression Omnibus (GEO: https://www.ncbi.nlm.nih.gov/geo/) under the accession number of GSE140911. ChIP-seq and RNA-seq data used from (32) GSE113335, and (41) GSE43231.

### Acknowledgements

We thank Monash University FACS core facility. We thank Xiaofeng Wang for Illumina HiSeq2500 high-throughput Sequencing at Harvard Medical School; as well as the Genewiz high-throughput sequencing facility, China.

## Author contributions

M.C.C., P.T., M.J.B., L.H.W. and P.P.D. performed experiments and analyzed the data. M.C.C., P.T., M.J.B., Y.K., S.J.T., S.L., L.P., S.H.O., and P.P.D. interpreted the data. M.C.C., Y.K., M.O., S.L., and L.P. performed bioinformatics analyses. M.C.C., P.T., S.H.O., and P.P.D. wrote the manuscript. All authors read and approved the final manuscript.

