## Supplemental material for "A saturating mutagenesis CRISPR-Cas9 mediated functional genomic screen identifies *cis-* and *trans-* regulatory elements of *Oct4* in murine ESCs"

Fig. S1

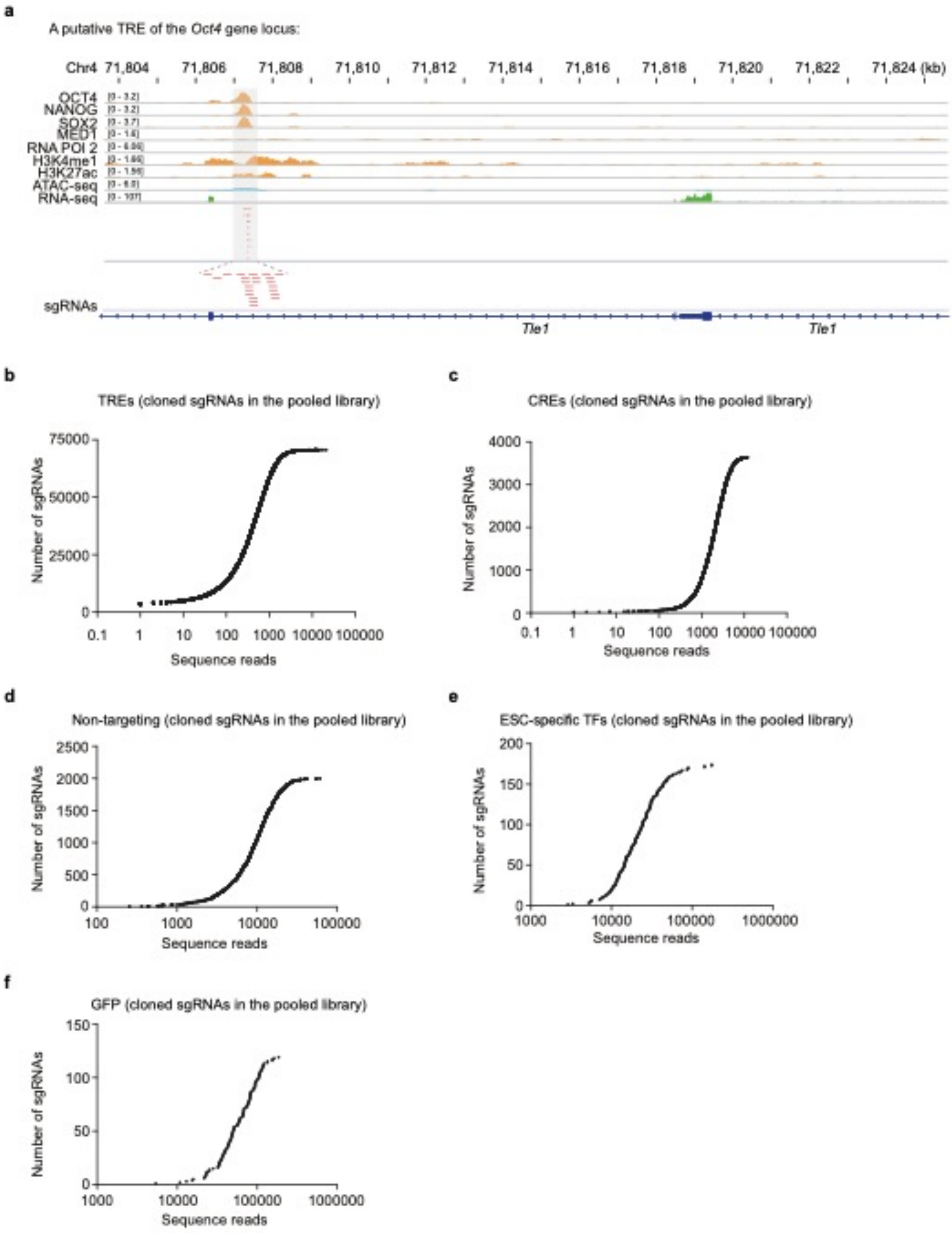

a
A putative TRE of the Oct4 gene locus:
Chr4 71,804 71,806 71,808 71,810 71,812 71,814 71,816 71,818 71,820 71,822 71,824 (kb)
OCT4
NANOG
SOX2
MED1
RNA POl 2
H3K4me1
H3K27ac
ATAC-seq
RNA-seq
sgRNAs
Tle1
Tle1
b
TREs (cloned sgRNAs in the pooled library)
Number of sgRNAs
Sequence reads
c
CREs (cloned sgRNAs in the pooled library)
Number of sgRNAs
Sequence reads
d
Non-targeting (cloned sgRNAs in the pooled library)
Number of sgRNAs
Sequence reads
e
ESC-specific TFs (cloned sgRNAs in the pooled library)
Number of sgRNAs
Sequence reads
f
GFP (cloned sgRNAs in the pooled library)
Number of sgRNAs
Sequence reads

**Figure S1. Design of a saturating CRISPR-Cas9 pooled library for ctSCAN-SMS.** (a) Genomic tracks show co-occupancy of ESC-TFs (OCT4, NANOG, SOX2), mediator (MED1), enhancer histone marks (H3K27ac, H3K4me1), and RNA Pol2 at a putative TRE of the *Oct4* gene locus in mESCs. ATAC-seq track represents open chromatin regions and enhancers (ENs); RNA-seq track represents gene expression. The highlighted region marked as a putative TRE. To target this putative TRE, we designed sgRNAs (shown with red dashed lines) upstream of all PAM sequences (NGG) within this TRE. (b-f) Representation of cloned sgRNAs in the REs CRISPR-Cas9 pooled library. The pooled library consisted of 74,576 sgRNAs in total that targeted CREs, TREs, GFP of the *Oct4*-GFP reporter, mESC-specific TFs, and non-targeting regions. All these sgRNAs were synthesized, pooled and cloned into a lentiviral vector; followed by deep-sequencing to ensure representation of the sgRNAs. Each dot represents an sgRNA and its corresponding read count in the pooled library. Y-axis: sgRNAs; X-axis: number of reads per sgRNA.

**Fig. S2**

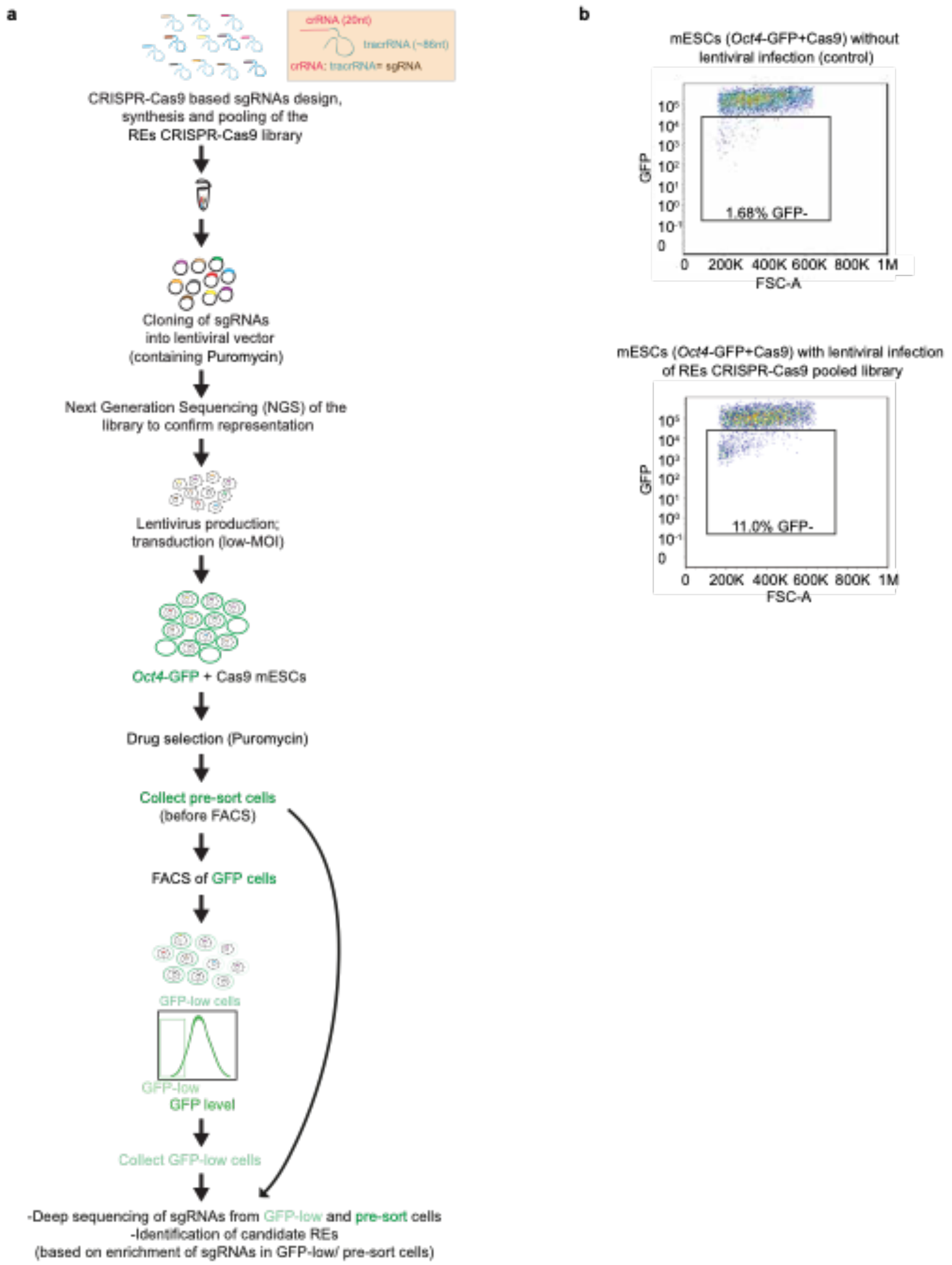
a
crRNA (20nt)
tracrRNA (~86nt)
crRNA: tracrRNA= sgRNA
CRISPR-Cas9 based sgRNAs design, synthesis and pooling of the REs CRISPR-Cas9 library
Cloning of sgRNAs into lentiviral vector (containing Puromycin)
Next Generation Sequencing (NGS) of the library to confirm representation
Lentivirus production; transduction (low-MOI)
Oct4-GFP + Cas9 mESCs
Drug selection (Puromycin)
Collect pre-sort cells (before FACS)
FACS of GFP cells
GFP-low cells
GFP-low
GFP level
Collect GFP-low cells
-Deep sequencing of sgRNAs from GFP-low and pre-sort cells
-Identification of candidate REs
(based on enrichment of sgRNAs in GFP-low/ pre-sort cells)
b
mESCs (Oct4-GFP+Cas9) without lentiviral infection (control)
GFP
FSC-A
1.68% GFP-
0
200K
400K
600K
800K
1M
mESCs (Oct4-GFP+Cas9) with lentiviral infection of REs CRISPR-Cas9 pooled library
11.0% GFP-

**Figure S2. Experimental design of ctSCAN-SMS to identify REs of the *Oct4* gene.** (a) A schematic diagram represents an outline of the CRISPR-Cas9 screen. (b) FACS analysis represents percentage (%) of GFP-negative (GFP-low) cells from the *Oct4*-GFP+Cas9 reporter mESCs either with no lentiviral infection (control) or with lentiviral infection of REs CRISPR-Cas9 pooled library at low multiplicity of infection (MOI).

**Fig. S3**

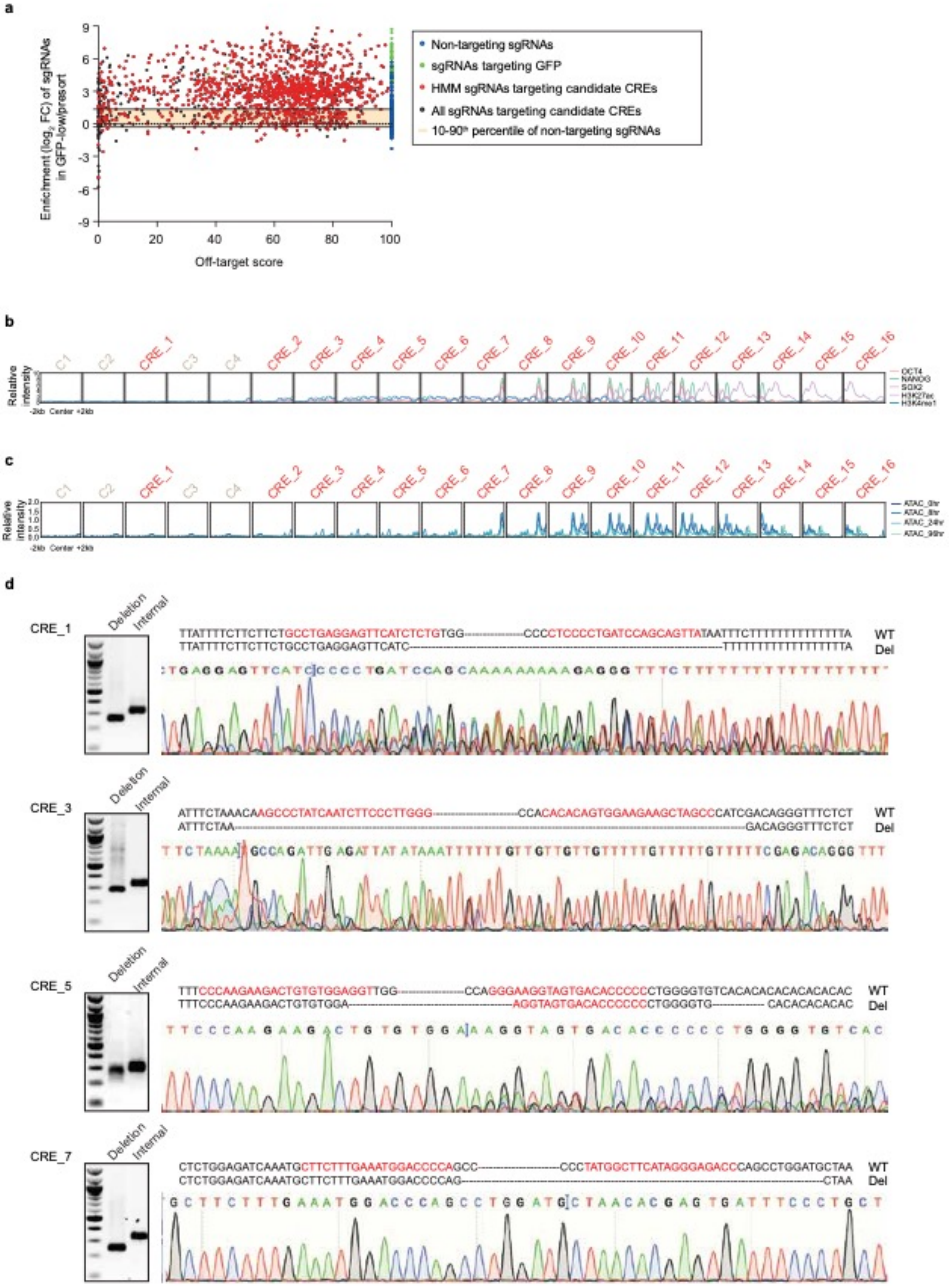
a
Enrichment ($log_2$ FC) of sgRNAs in GFP-low/presort
Off-target score
Non-targeting sgRNAs
sgRNAs targeting GFP
HMM sgRNAs targeting candidate CREs
All sgRNAs targeting candidate CREs
10-90th percentile of non-targeting sgRNAs
b
C1
C2
CRE_1
C3
C4
CRE_2
CRE_3
CRE_4
CRE_5
CRE_6
CRE_7
CRE_8
CRE_9
CRE_10
CRE_11
CRE_12
CRE_13
CRE_14
CRE_15
CRE_16
Relative intensity
-2kb Center +2kb
c
Relative intensity
-2kb Center +2kb
d
CRE_1
CRE_3
CRE_5
CRE_7
Deletion
Internal
WT
Del

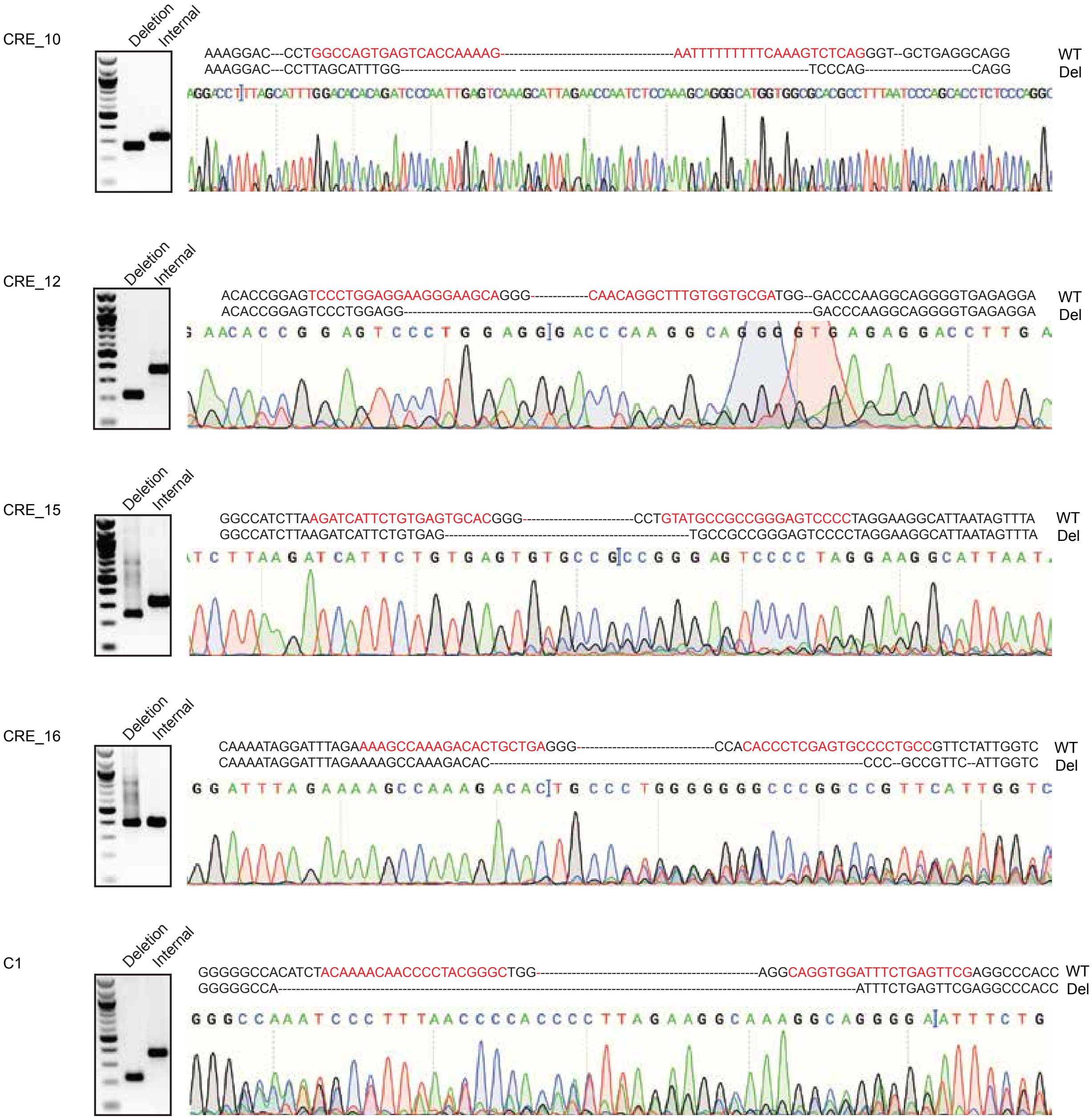


**Figure S3. Identification of candidate CREs of the *Oct4* and their regulatory features.** (a) Dot plot analysis shows the enrichment score of each sgRNA by comparing their frequency in the GFP-low cells to the pre-sort cells. Off-target scores were calculated for each sgRNA and ranked from left to right (lowest to highest off-target scores) on the x-axis. A higher off-target score signifies fewer off-targets for a particular sgRNA; off-target score >10 was used as a cut off for inclusion in analyses. Hidden Markov model (HMM) analysis representing functionally significant sgRNAs. sgRNAs targeting GFP and non-targeting sgRNAs were used as positive and negative controls, respectively. The shaded region highlights the $10^{th}$-$90^{th}$ percentiles of non-targeting sgRNAs, and other sgRNAs targeting CREs. (b) Profile plots display occupancy changes of OCT4, NANOG, SOX2, H3K27ac, H3K4me1 within the CREs of *Oct4*. (c) ATAC-seq represents the changes of open chromatin regions/ENs from undifferentiated to the differentiated mESC state (0, 8, 24 and 96 hr) within the CREs of *Oct4*. (d) Paired sgRNAs-mCherry (5' and 3' sgRNAs) were introduced to target both the flanking ends of each CRE to create a deletion in wild-type mESCs. Two PCRs (deletion PCR = to detect deleted allele; internal PCR = to detect wild-type allele) were used for each targeted CRE, which show "bull/mixed" population of edited cells from mCherry-positive cells. Deleted PCR products were excised, and Sanger sequencing was performed to confirm the targeted CRE deletions. sgRNA sequences are highlighted in red. Deletions of few active CREs are shown here. Deletion sizes, sgRNAs and genotyping primers are listed in Table S3.

### Fig. S4

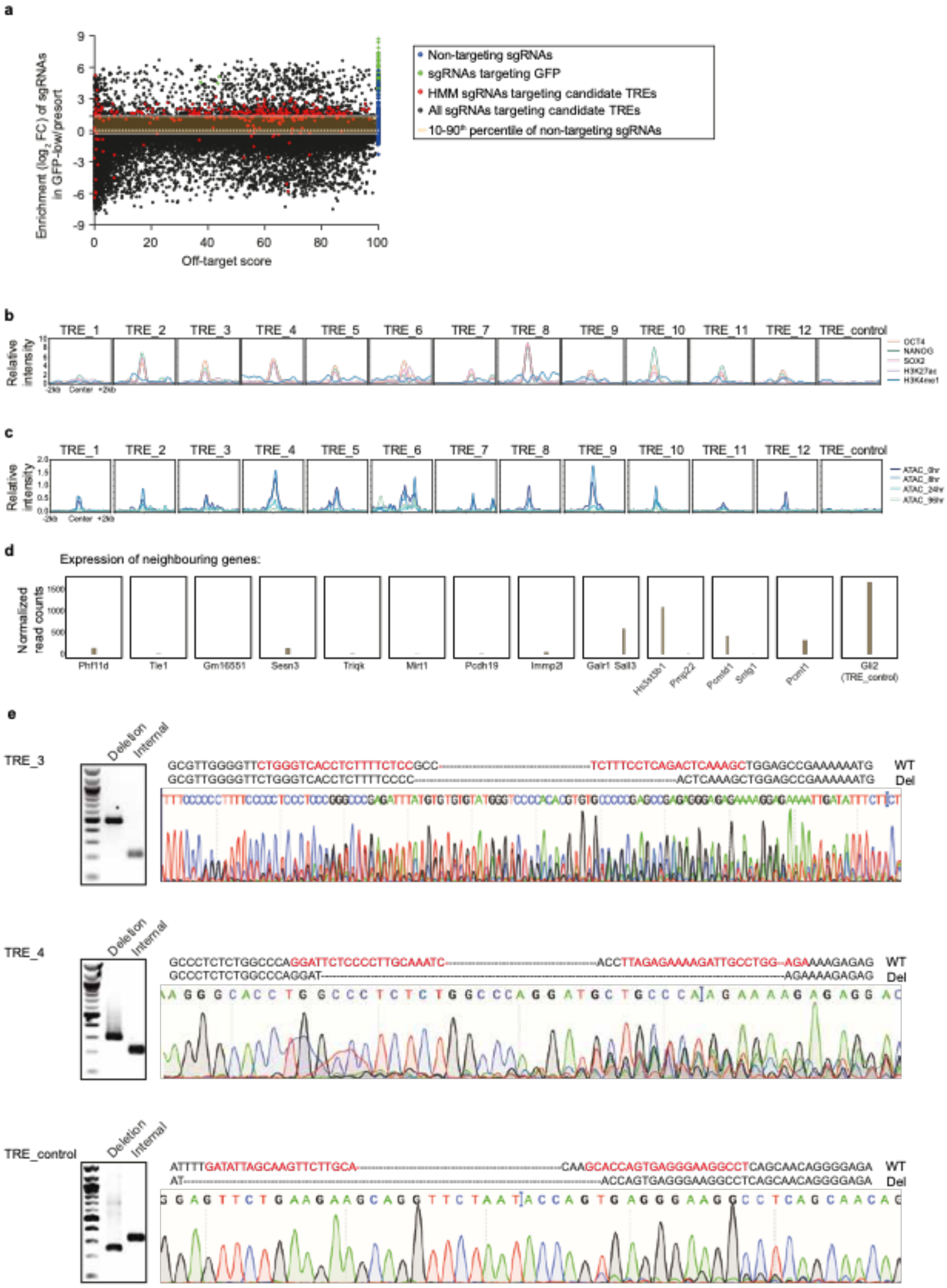
a
Enrichment ($log_2$ FC) of sgRNAs in GFP-low/presort
Off-target score
Non-targeting sgRNAs
sgRNAs targeting GFP
HMM sgRNAs targeting candidate TREs
All sgRNAs targeting candidate TREs
10-90th percentile of non-targeting sgRNAs
b
TRE_1 TRE_2 TRE_3 TRE_4 TRE_5 TRE_6 TRE_7 TRE_8 TRE_9 TRE_10 TRE_11 TRE_12 TRE_control
Relative intensity
OCT4
NANOG
SOX2
H3K27ac
H3K4me1
-2kb Center +2kb
c
TRE_1 TRE_2 TRE_3 TRE_4 TRE_5 TRE_6 TRE_7 TRE_8 TRE_9 TRE_10 TRE_11 TRE_12 TRE_control
Relative intensity
ATAC_0hr
ATAC_8hr
ATAC_24hr
ATAC_96hr
-2kb Center +2kb
d
Expression of neighbouring genes:
Normalized read counts
Phf11d Tle1 Gm16551 Sesn3 Triqk Mirt1 Pcdh19 Immp2l Galr1 Sall3 Hs3st3b1 Pmp22 Pcmtd1 Sntg1 Pcnt1 Gli2 (TRE_control)
e
TRE_3
Deletion Internal
GCGTTGGGGTTCTGGGTCACCTCTTTTCTCCGCC---TCTTTCCTCAGACTCAAAGCTGGAGCCGAAAAAATG WT
GCGTTGGGGTTCTGGGTCACCTCTTTTCCCC---ACTCAAAGCTGGAGCCGAAAAAATG Del
TRE_4
Deletion Internal
GCCCTCTCTGGCCCAGGATTCTCCCCTTGCAAATC---ACCTTAGAGAAAAGATTGCCTGG--AGAAAAGAGAG WT
GCCCTCTCTGGCCCAGGAT---AGAAAAGAGAG Del
TRE_control
Deletion Internal
ATTTTGATATTAGCAAGTTCTTGCA---CAAGCACCAGTGAGGGAAGGCCTCAGCAACAGGGGAGA WT
AT---ACCAGTGAGGGAAGGCCTCAGCAACAGGGGAGA Del

**Figure S4. Identification of candidate TREs of the *Oct4* and their regulatory features.** (a) Dot plot analysis exhibits the enrichment score of each sgRNA. Off-target scores were calculated for each sgRNA and ranked them left to right (with lowest to highest off-target scores) on the x-axis. A higher off-target score signifies fewer off-targets for a particular sgRNA; off-target score >10 was used as a cut off for inclusion in analyses. Hidden Markov model (HMM) analysis representing functionally significant sgRNAs. sgRNAs targeting GFP and non-targeting sgRNAs were used as positive and negative controls, respectively. The shaded region highlights $10^{th}$-$90^{th}$ percentiles of non-targeting sgRNAs and other sgRNAs that target TREs. (b) Binding profile of OCT4, NANOG, SOX2, H3K27ac, H3K4me1 across the different TREs of *Oct4.* (c) Open chromatin regions/ENs changes within the TREs captured by ATAC-seq at different time points from undifferentiated to the differentiated mESC state (0, 8, 24 and 96hr). (d) RNA-seq data represents the expression of neighbouring genes of the corresponding TREs. (e) Paired sgRNAs-mCherry (5' and 3' sgRNAs) were introduced to target both the flanking ends of each TRE to create a deletion in wild-type mESCs. Two PCRs (deletion PCR = to detect deleted allele; internal PCR = to detect wild-type allele) were used for each targeted TRE, which show "bull/mixed" population of edited cells from mCherry-positive cells. Deleted PCR products were excised, and Sanger sequencing was performed to confirm the targeted TRE deletions. sgRNA sequences are highlighted in red. Deletions of only two active TREs are shown here. Deletion sizes, sgRNAs and genotyping primers are listed in Table S3.

**Fig. S5**

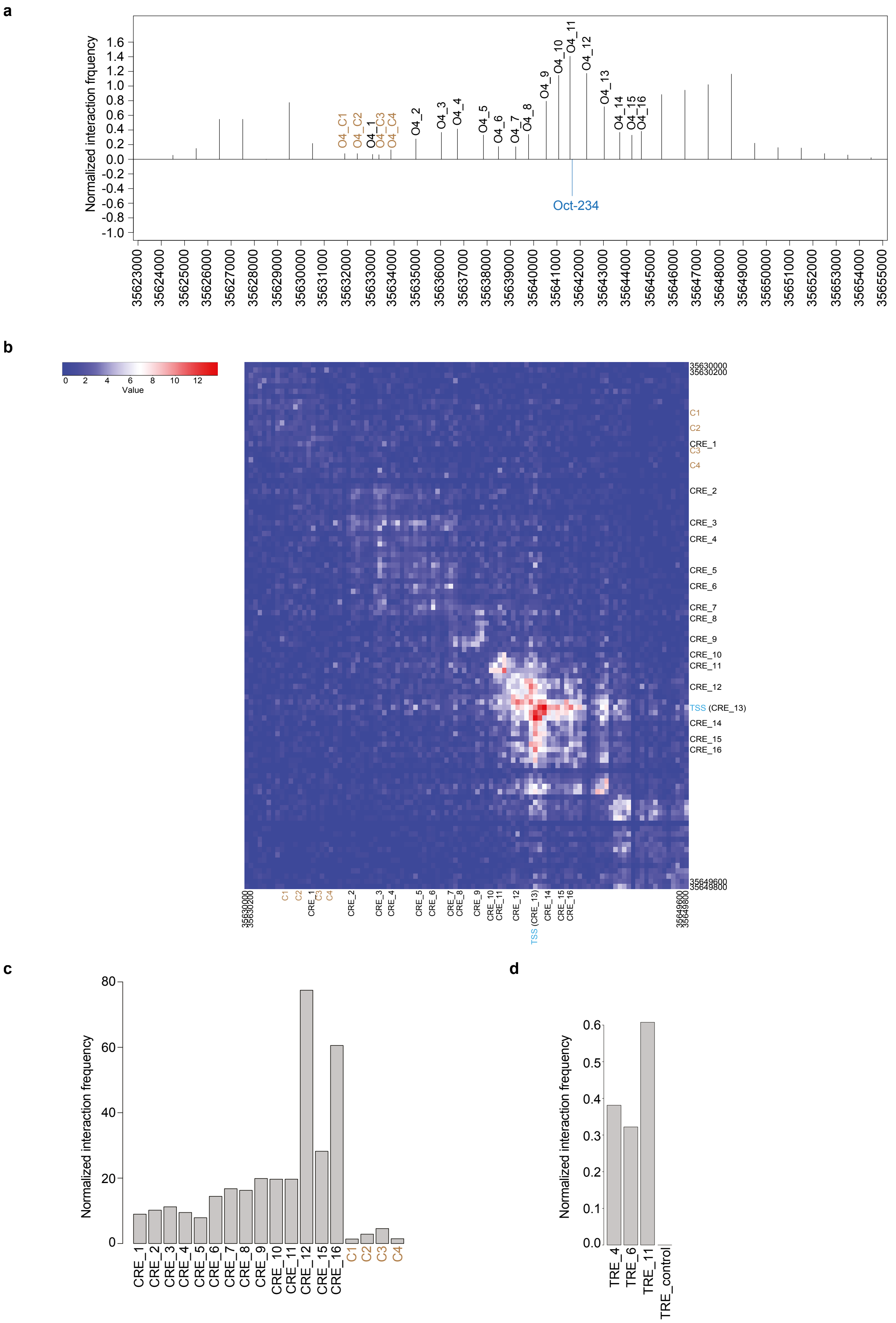
a
Normalized interaction frquency
O4_C1
O4_C2
O4_1
O4_C3
O4_C4
O4_2
O4_3
O4_4
O4_5
O4_6
O4_7
O4_8
O4_9
O4_10
O4_11
O4_12
O4_13
O4_14
O4_15
O4_16
Oct-234
b
Value
C1
C2
CRE_1
C3
C4
CRE_2
CRE_3
CRE_4
CRE_5
CRE_6
CRE_7
CRE_8
CRE_9
CRE_10
CRE_11
CRE_12
TSS (CRE_13)
CRE_14
CRE_15
CRE_16
c
Normalized interaction frequency
d
Normalized interaction frequency
TRE_4
TRE_6
TRE_11
TRE_control

**Figure S5. Physical interactions between CREs, TREs and the *Oct4* promoter.** (a) 4C-seq data representing normalized interaction frequencies between CREs and the *Oct4* promoter. Oct4-234, a region ~1.5 kb upstream of TSS of the *Oct4*, was used as a “viewpoint” to analyse the contact frequencies between the viewpoint and CREs. The normalized interaction/contact frequencies between CREs and the *Oct4* promoter was measured at a 1kb resolution window. (b) Micro-C data illustrating the contact matrix of all intra-chromosomal interactions/contacts between any two genomic loci around ~20kb region (35,630,000 to 35,650,000) of the *Oct4* gene locus (TSS is at ~ 35,643,000) at a 200bp resolution. Scale represent lowest (blue) – to intermediate (white) – to highest (red) contact frequencies. (c) Bar plot displaying (from Micro-C data) normalized interaction frequencies between CREs and the *Oct4* promoter at a 200bp resolution. (d) Bar plot showing (from Micro-C data) normalized interaction frequencies between TREs and the *Oct4* promoter at a 1kb resolution.

**Fig. S6**

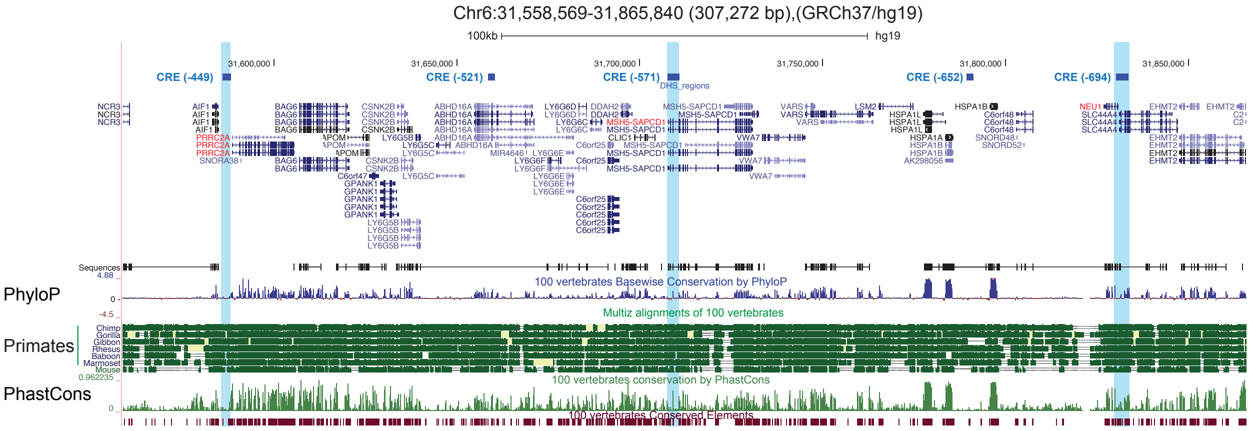


**Figure S6. Conserved human active CREs of *OCT4.*** Orthologous sequences of three highlighted high-confidence CREs (-449, -571, -694) of human *OCT*4 are shown among representative primates and mouse. PhyloP and PhastCons estimate evolutionary conservation among 100 vertebrates.
